# Microbe-mediated plant acclimation to drought may be rare in agriculture

**DOI:** 10.64898/2026.04.02.715620

**Authors:** Mia M. Howard, Lana G. Bolin, Glade D. Bogar, Sarah E. Evans, Jay T. Lennon, Sandra T. Marquart-Pyatt, Jennifer A. Lau

## Abstract

Microbial communities can shift under drought in ways that enhance plant performance during drought (“microbe-mediated acclimation”). However, it is also possible for microbial communities to shift in ways that worsen the effects of drought (“mal-acclimation”). It is unclear how and where microbe-mediated acclimation vs. mal-acclimation occurs, or if there are types of soils or microbial communities that are more likely to harbor microbes that enhance plant acclimation and limit mal-acclimation. We tested for microbe-mediated plant acclimation/mal-acclimation to drought in soils from 21 maize farms in the midwestern United States, spanning a range of climate, soil types, and management practices. We first conditioned soil microbial communities to drought vs. well-watered conditions in a greenhouse and then tested for microbe-mediated acclimation by growing maize in soils inoculated with the conditioned microbial communities under drought and well-watered conditions. Drought-conditioned soils did not enhance plant performance under drought. In fact, one third of the farms exhibited mal-acclimation, especially under well-watered conditions where wet-conditioned soils reduced plant performance in well-watered contemporary conditions. Farm management practices, climate, soil texture, and microbial diversity generally did not predict when this microbe-mediated mal-acclimation occurred. Overall, these results suggest that in agricultural soils, microbes may frequently impede–rather than facilitate–plant acclimation to soil moisture levels.

**Open research statement:** The plant and soil data used in this study are available via the Environmental Data Initiative repository at https://doi.org/10.6073/pasta/f4a0db3a076cf6d8cef908947f82736e. The bacterial and fungal amplicon sequence data are available via the European Nucleotide Archive under accessions PRJEB110071 and PRJEB109827, respectively.

## INTRODUCTIONS

Microbiomes consist of many diverse species of bacteria, fungi, and archaea that have the potential to influence host nutrition, health, and performance. The surge in microbiome research over the past two decades highlighted the importance of microbiomes for host biology and has sparked interest in how microbes could be used to improve the health of plants and animals and how microbes might facilitate the acclimation of their hosts to rapidly changing environments. Yet, microbes are not always beneficial, and some taxa or communities reduce host fitness and can exacerbate stressful conditions for their hosts.

With an increasing frequency and severity of droughts (Chiang et al. 2021), there is a particular need to understand how microorganisms influence plant resilience to water stress. Microbes can improve plant performance during drought through altering both their host’s physiology [e.g., synthesizing antioxidants (Hamilton and Bauerle 2012) and phytohormones (Khan et al. 2012)] and environment [e.g., by producing biofilms that retain water in the rhizosphere (Yang et al. 2021) and promote soil aggregation (Vardharajula and Z 2014; Prasanna Kumar et al. 2022)]. Some microbes such as mycorrhizal fungi can also extend the reach of plant roots, helping them access distant sources of water, or improve nutrient availability under stressful conditions (Gehring et al. 2017). Many studies have investigated the physiological mechanisms through which microbes can enhance drought tolerance (Ngumbi and Kloepper 2016), but microbes can also intensify the negative consequences of drought stress for their plant hosts. For example, pathogens can be more damaging when hosts are drought-stressed and more susceptible to infection (Choudhary and Senthil-Kumar 2024). As a result, it remains unclear how often the diverse microbial communities found in the soil promote vs. hinder plant acclimation to drought.

In some cases, microbial communities shift compositionally under stress in ways that increase the stress tolerance benefits they provide to their hosts. “Microbe-mediated acclimation” occurs when the exposure of microbes to a stress enhances the performance of plants under that stress condition compared to microbes to that stress [Bolin 2023; see also Mueller et al. 2020; Trivedi et al. 2020, and “microbe-mediated adaptation” in Petipas et al. (2021)]. For example, in the context of soil moisture, microbe-mediated acclimation could occur if drought increased the relative abundance of drought-tolerant biofilm producers in a soil microbial community, increasing the retention of water in the soil and buffering the effects of drought stress on plants (Bolin et al. 2023). Likewise, microbe-mediated acclimation would occur if drought reduced the relative abundance (or activity) of pathogens in the soil, resulting in improved plant survival under drought where pathogen infection may be more detrimental. Microbe-mediated acclimation to high soil moisture could also occur if microbes from wet soils enhanced plant performance in wet conditions. Alternatively, “microbe-mediated *mal-acclimation*” occurs when the microbial communities that develop under a certain condition exacerbate the negative effects of that condition on their hosts. For example, microbe-mediated mal-acclimation to drought occurs when the microbial communities exposed to drought worsen the effects of drought on plants. This could occur, for example, if drought reduces the amount of carbon that plants release into the rhizosphere through root exudates to support beneficial bacteria or mycorrhizal fungi (Gehring et al. 2017) that might be particularly helpful in drought-stressed soils where nutrients are less mobile (Suriyagoda et al. 2014; Nguyen et al. 2018). Microbe-mediated mal-acclimation could also stem from the accumulation of pathogens in the soil–particularly pathogens that are adapted to the stress condition. Pathogens adapted to high moisture soils, for example, could drive microbe-mediated malacclimation to wet conditions. Regardless of the mechanism, microbial modulation of plant responses to soil moisture could play an important role in the resilience of plant communities in natural and managed landscapes.

Over the past decade, a growing number of studies suggest that microbes frequently help plants acclimate to stresses such as drought. Researchers have found evidence for microbe-mediated acclimation to drought across diverse plant species (Table 1). In each of these cases, plant acclimation to drought stress is determined at least in part by the response of associated microbial communities to drought. In total, these examples suggest that microbial communities frequently respond to drought stress in ways that promote plant acclimation to drought, but the relatively few studies to date mean that the prevalence of this phenomenon remains to be determined. Studies that tested multiple plant species indicate that the extent of microbe-mediated acclimation can vary by plant species. For example, microbes from drier sites improved the performance of the grass *Lagurus ovatus* and the forb *Sisymbrium erysimoides* under drought stress, but not the legume *Trifolium stellatum* (O’Brien et al. 2018). Moreover, studies that experimentally evaluate microbe-mediated acclimation typically only test a single microbial community (i.e., soil collected from a single location), which researchers experimentally condition under drought versus well-watered conditions in a controlled environment (Lau and Lennon 2012; Ricks and Yannarell 2023, see Table 1). However, soil microbial communities can vary functionally between habitats, even between habitats of the same type (Bauer et al. 2017), and even within the same site, including in relatively homogeneous environments such as conventional maize fields (Suriyavirun et al. 2019). Thus, soil communities are likely to vary widely in the degree to which they change under drought and harbor drought-resistant microorganisms that could improve plant performance under drought.

**Table 1.** Evidence for microbe-mediated acclimation to drought.

| Plant species | Microbe source | Summary | Citation |
| --- | --- | --- | --- |
| *Brassica rapa*<br>(Brassicaceae) | Early successional grassland soil microbial communities conditioned under wet and dry conditions in greenhouse | Conditioned soil microbes in a greenhouse under wet vs. dry soil moisture. Microbes improved the performance of plants under their acclimated condition. | (Lau and Lennon 2012) |
| *Brassica rapa*<br>(Brassicaceae) | Tall-grass prairie soil microbial communities conditioned under wet and dry conditions in greenhouse | Conditioned soil microbes in a greenhouse under wet vs dry soil moisture, as well as with or without a plant. Microbes improved the performance of plants under their conditioned environment, regardless of whether a plant was present in the conditioning stage | (Ricks and Yannarell 2023) |
| *Panicum virgatum*<br>(Poaceae) | Endophytes isolated from grasses across a natural precipitation gradient | Endophytic fungi that were more tolerant to water stress tended to help their host plants under drought stress, and isolates from drier sites were generally more beneficial to plants | (Giauque et al. 2019) |
| *Ostrya virginiana*<br>(Betulaceae),<br>*Betula nigra*<br>(Betulaceae) | Sites across a natural precipitation gradient | Microbes from drier sites improved plant performance under drier conditions | (Allsup and Lankau 2019) |
| *Trifolium stellatum*<br>(Fabaceae),<br>*Lagurus ovatus*<br>(Poaceae),<br>*Sisymbrium erysimoides*<br>(Brassicaceae) | Sites across a natural precipitation gradient (soils collected underneath Retama shrubs) | Microbes from drier sites improved performance (biomass) of the grass and crucifer under drier conditions, but not the legume | (O'Brien et al. 2018) |

Furthermore, it is unclear what characteristics of soils or their microbial communities enhance plant acclimation to stress. Larger or more diverse communities might be more functionally resilient to stressors such as drought or more likely to contain plant beneficial microbes that can also tolerate drought (i.e., the sampling/portfolio effect (Philippot et al. 2021)). Therefore, these communities might be more likely to lead to microbe-mediated acclimation. Sites that frequently experience drought–or have experienced drought more recently–might contain more drought-tolerant microbes than wetter sites and thus be more stable under drought stress (Dacal et al. 2022) and quicker to influence plants (either negatively or positively) under drought. Understanding the characteristics of soils and microbial communities that promote microbe-mediated acclimation will help us predict where and when it may occur.

In this study we investigated whether microbe-mediated acclimation to drought occurs across a broad range of soil microbial communities, focusing on maize, a dominant agricultural crop. Large-scale maize agriculture presents an opportunity to assess the prevalence of microbe-mediated drought acclimation in soils across a geographic range encompassing different soil types and climatic variation in both temperature and precipitation, as well as variation in management (i.e., tillage, irrigation). To assess the prevalence of microbe-mediated acclimation and mal-acclimation, we tested soil microbial communities from maize farms in the midwestern United States for their capacity to respond to drought in ways that promote plant acclimation to drought (i.e., microbe-mediated acclimation to drought), and tested whether soil, microbial community, climate, and management characteristics predicted these effects.

## METHODS

### Study design

We tested whether soil microbial communities from 21 midwestern maize farms promote microbe-mediated acclimation or mal-acclimation to drought in a two-phase experiment (Fig 1A). First, in the “conditioning phase”, we conditioned each soil community under drought or well-watered conditions for one (maize) plant generation to generate inocula with different “microbe histories”. Then, in the “drought response phase”, we inoculated maize plants with the conditioned microbial communities and grew them under drought and well-watered conditions in a full factorial design and assessed plant performance. Microbe-mediated acclimation to drought occurs when microbes previously exposed to drought improve plant performance under contemporary drought (Fig. 1B). Conversely, microbe-mediated mal-acclimation occurs when microbes previously exposed to drought reduce plant performance under contemporary drought (Fig. 1C). Likewise, microbe-mediated acclimation and mal-acclimation to well-watered conditions occurs when microbes exposed to well-watered conditions enhance (Fig. 1B) or reduce (Fig. 1C) plant performance under contemporary well-watered conditions, respectively.

**Fig. 1.**
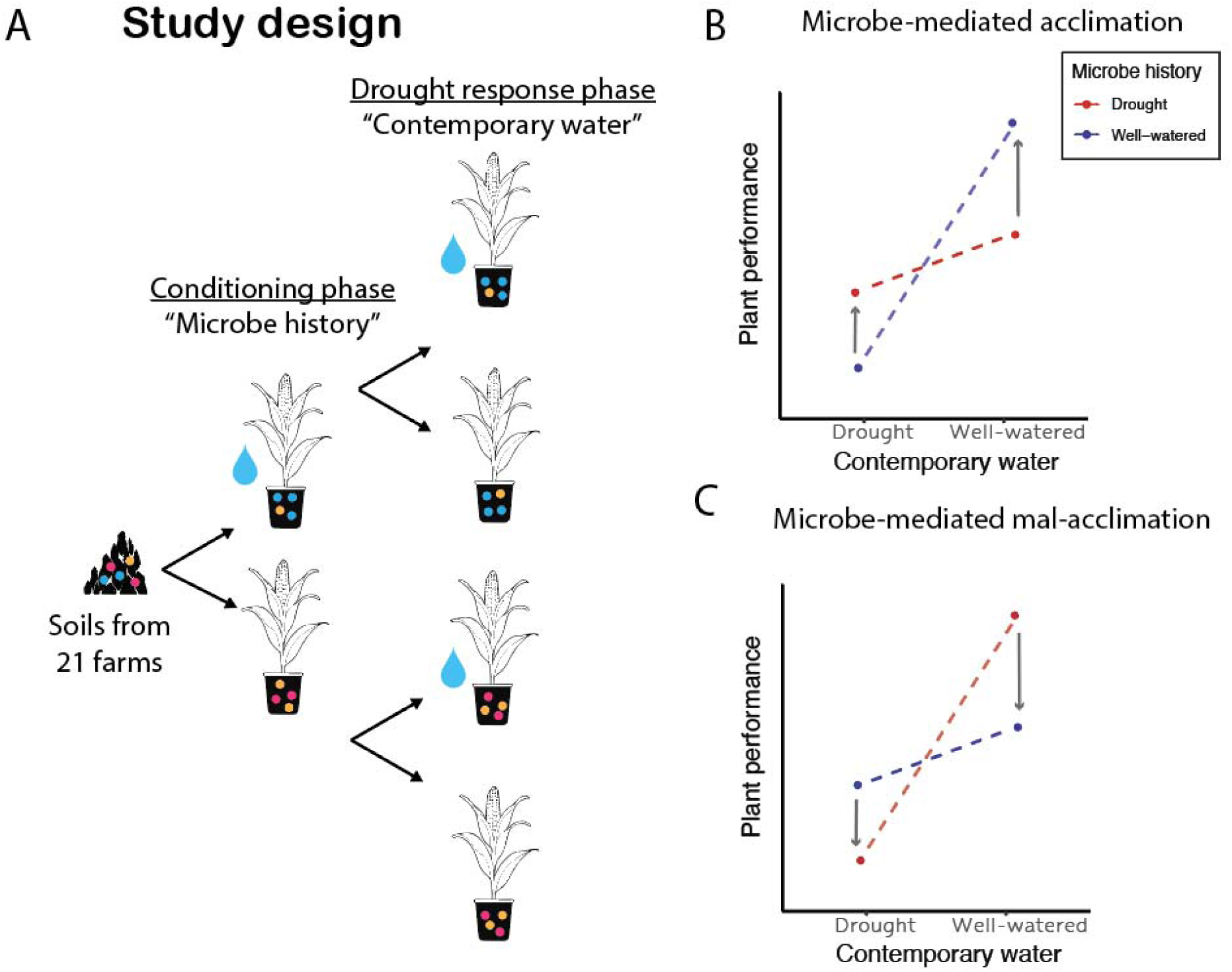
Design of experimental conditioning of soil microbes to drought (“microbe history”) and subsequent assessment of microbe-mediated plant responses to drought (“contemporary water”) (A). (B) Microbe-mediated acclimation occurs when drought-conditioned microbes improve plant performance under drought, and/or when wet-conditioned microbes improve plant performance under well-watered conditions (this plot illustrates microbe-mediated acclimation in both watering conditions). (C) Microbe-mediated mal-acclimation occurs when microbes reduce plant performance in the watering environment under which they were conditioned. Arrows indicate the relative benefit (in B) or detriment (in C) of the history of the inoculant matching contemporary watering conditions. These microbe-mediated acclimation and mal-acclimation effects are characterized by an interactive effect between microbe history and contemporary conditions.

### Source inocula

We sourced soil communities from 68 maize fields across Illinois, Indiana, and Michigan in September and October, 2021. Farms were identified from a large-scale farmer survey (Panel Farmer Survey, (Guo et al. 2023; Irvine et al. 2023)) and chosen to balance a range of management approaches (e.g., irrigation, tillage) and geography. Soils were sampled to a depth of 20 cm, with 15 subsamples taken in a straight line (one every 2 paces) parallel to the field edge, from a randomly-selected starting point approximately 10 m (15 paces) into the field. Soils were transported on ice to one of two labs (at Kellogg Biological Station, Hickory Corners, MI or Indiana University, Bloomington, IN) and stored at 4 °C until homogenization by sieving to 4 mm, followed by subsampling into air-dried, field-moist, and frozen (−20 °C) portions for further analysis (described in later sections). Duration of storage at 4 °C prior to sieving ranged from 1 to 12 d (mean 4.8).

### Soil conditioning phase (“microbe history”)

To condition soil microbial communities under different soil moisture conditions, we inoculated pots of steam-pasteurized potting media with each field-collected soil community, planted maize seeds in them, and grew the plants in either well-watered or drought conditions. Specifically, we inoculated 5 L pots with each soil microbial community at a rate of 4% (v/v) onto potting media designed to simulate the agricultural field soils observed across our sites [equal parts by volume sand, calcined clay (Turface MVP, Buffalo Grove, IL, USA), and MetroMix820 (SunGro Horticulture, Agawam, MA, USA), and natural field topsoil from Bloomington, IN, USA]. To reduce the existing microbial community in the background potting media, we steam-pasteurized the media twice for 4 h each time with a 24 h rest period in between. We inoculated the mesocosms on October 26, 2021, and planted two maize seeds (Pioneer PO306CYFR, Pioneer, Johnston, IA, USA), thinning to one seedling upon germination. We replicated each of the 69 farm × watering treatment (well-watered or drought) combinations six times for a total of 828 mesocosms, plus 30 uninoculated plants. The experiment took place in the greenhouses at Indiana University, with the pots arranged in a randomized order. We began the watering treatments 17 d after planting. The well-watered group was watered as needed to avoid water stress (typically daily), while the drought group was only watered when some plants were visibly wilting (i.e., drooping, curling leaves). We harvested the maize after 17 weeks and collected the conditioned soils from the mesocosms (now with well-watered or drought “microbe histories”) to inoculate plants in the drought response phase (see below). We selected 21 representative soil microbial communities out of the initial pool of 68 fields to test for microbe-mediated acclimation in our study. These communities were selected based on variation in their effects on plant drought responses and representation across geography and management practices. Of the 21 farms we ultimately tested for microbe-mediated acclimation to drought, six were in Illinois, seven were in Indiana, and six were in Michigan. In terms of management, 38% used irrigation, while the remainder relied only on natural rainfall. In terms of tillage, 14% used conventional tillage practices, 57% practiced conservation tillage, and 29% were no-till. We stored the conditioned soils at 4 °C between the soil conditioning and drought response phases.

### Drought response phase (“contemporary water”)

To test if microbes responded to drought in ways that benefit plants under drought (i.e., to test for microbe-mediated acclimation), we inoculated plants with microbes conditioned in well-watered or drought conditions (“microbe history”) and grew them under well-watered or drought conditions in a greenhouse. We inoculated six replicate pots per inoculant per contemporary watering condition; with three inoculants per farm per conditioning water treatment, this yielded a total of 756 pots, which were split into two contemporary watering treatments (drought and well-watered). We transferred our conditioned microbes to 5 L pots of pasteurized potting media (due to the large volume of soil needed, we used the same media from the conditioning phase after steam pasteurizing it twice using the same methods) at 10% (v/v) on April 7, 2022, and planted two maize seeds per pot (later thinned to one) the following day. We started the watering treatments 17 d after planting using the same watering protocols as in the conditioning phase and fertilized (Peter’s 20-20-20, Everris NA, Inc., Dublin, OH, USA) 31, 49, and 65 d after planting. Once plants began to flower (67 d after planting), we hand-pollinated flowering individuals every other day to promote kernel filling until flowering stopped.

### Plant performance measurements

During the drought response phase, we recorded plant height, number of collared leaves, and stem diameter at 59 d after planting. At harvest (104 d after planting), we collected and dried aboveground plant biomass and counted ears produced. We weighed vegetative mass, and reproductive biomass as complete ears, cobs (ears with the husks and silks removed), and kernels. Due to asynchronous flowering and imperfect pollination in the greenhouse (an average cob was only 56% covered with kernels), we also estimated kernel mass for each plant under complete kernel set. We calculated this “scaled kernel mass” by visually estimating the proportion of kernel set for each cob and multiplying measured kernel mass by the inverse of the proportion of kernel set.

### Data analysis

To test for microbe-mediated acclimation, we used linear mixed effects models with the plant performance response variables of height, number of collared leaves, vegetative biomass, ear mass, cob mass, kernel mass, or scaled kernel mass; with fixed effects of microbe history, contemporary water treatment, and their interaction; and a random effect of inoculant replicate (nested within farm and microbe history). We analyzed ear number using Poisson regression with the same effects structure. A significant microbe history × contemporary water interaction indicates that microbes’ historical soil moisture conditions affect plant responses to current soil moisture. We assessed statistical significance after adjusting p-values with a false discovery rate correction (FDR) to account for comparison across multiple measures of plant performance (Benjamini and Hochberg 1995). Microbe-mediated acclimation occurs when plants perform best when inoculated with microbes from a history that matches contemporary growth conditions (Fig. 1B), while microbe-mediated mal-acclimation occurs when microbes reduce plant performance under acclimated conditions (Fig. 1C). Main effects of microbe history indicate that the microbial communities have shifted in response to the conditioning phase treatments in ways that affect plant growth, but do not necessarily indicate microbe-mediated (mal)acclimation. For example, if a microbial history of drought improves plant performance under both well-watered and drought conditions, this would indicate that drought generally selects for beneficial microbes, but not microbes that specifically facilitate drought acclimation.

To test whether microbe-mediated acclimation varied by farm, we conducted similar analyses but included farm (and all interactions with farm) as a fixed, rather than random, factor. A significant microbe history × contemporary water × farm interaction indicates that microbe mediated (mal-)acclimation varies among farms. Because we detected this significant three-way interaction for the plant height response variable, we then analyzed each farm individually (21 separate models) for microbe-mediated acclimation using linear mixed effects models with plant height as the response variable; with microbe history, contemporary water treatment, and their interaction as fixed effects; and with inoculant replicate as a random effect.

We analyzed soil, microbial, climate, and management characteristics that predicted microbe-mediated acclimative responses in two different ways: individually and aggregated into principal component analysis (PCA) axes. Individual characteristics we tested were: soil bacterial and fungal diversity (Shannon diversity index), fungal pathogen load (based on relative abundance of putative plant pathogens), soil mycorrhizal fungi relative abundance, soil pH (2:1 field-moist 0.01M CaCl2), soil texture (% sand), microbial biomass (based on C), soil extracellular polymeric substance (EPS) content, soil wet aggregate stability, farm irrigation (irrigated or non-irrigated), farm tillage (no-till, conservation tillage, or conventional tillage), mean annual precipitation and mean annual temperature (30-year averages for each farm site). We fit linear models for each predictor variable that included the coefficient of the microbe history × contemporary water interaction term from models of plant height (β_microbe history × contemporary water_) as the response variable, and the farm characteristic as the predictor variable. A significant effect of farm characteristic would indicate that it predicted the microbe-mediated acclimation response. The climatic data were 30-year averages (1999-2020) from the PRISM database (PRISM Group). Soil and chemical characteristics, microbial community properties, and analysis methods and citations are described in Appendix 1, Section S1: Methods.

Because many other soil and microbial properties could also affect microbe-mediated acclimation, and because many soil and microbial properties covary, we additionally aggregated a wider array of factors into PCA axes for use as predictor variables using vegan (Oksanen et al., 2022). These included: 1) soil data [sand %, silt %, clay %, cation exchange capacity, air-dry pH, organic matter %, wet aggregate stability, and concentrations of plant nutrients (C, N, P, Ca, Mg, K, Na, B, Fe, Mn, Cu, Zn,)], and 2: microbial data [soil microbial biomass, EPS, Shannon diversity of bacteria and fungi, and relative abundance of fungal functional groups related to plants and soil (arbuscular mycorrhizal fungi, foliar endophytes, root endophytes, plant pathogens, root pathogens, leaf/fruit/seed pathogens, root associates, soil saprotrophs, litter saprotrophs, nectar saprotrophs, wood saprotrophs, dung saprotrophs, unspecified saprotrophs, mycoparasites, algal parasites)]. We then tested whether soil and microbial properties predicted microbe-mediated acclimation/mal-acclimation by fitting a linear model with β_microbe history× contemporary water_ as the response variable and the first two principal components of the soil properties PCA and the first two principal components of the microbial PCA as predictors. We performed all analyses using R version 4.2.2 (R Core Team 2022), fit mixed models using lme4 (Bates et al., 2015), and analyzed the models using car (Fox and Weinberg, 2019) and emmeans (Lenth, 2022).

## RESULTS

### Do agricultural soils typically promote microbe-mediated plant acclimation to drought?

With a few exceptions, soil microbial community exposure to drought stress worsened the negative effects of drought on plants, and soil microbial community exposure to wetter soil environments reduced plant growth in subsequent wet conditions (microbe history × contemporary moisture: all P < 0.05, Fig. 2, Appendix 1: Table S1), consistent with microbe-mediated mal-acclimation (Fig. 1C). Specifically, contemporary drought reduced all metrics of plant growth and reproduction except for number of ears (Fig. 2, Appendix S1: Table S1). Inoculating plants with drought-conditioned microbes generally further reduced maize growth and reproduction under drought. Likewise, microbes conditioned under ample soil moisture reduced plant performance under well-watered conditions. This general pattern of microbe-mediated mal-acclimation was observed for all metrics of both vegetative (Fig. 2A-C) and reproductive (Fig. 2D-H) performance traits with two exceptions. First, for biomass, drought-exposed microbial communities supported higher biomass under drought, but also under wet conditions, suggesting these microbial communities are generally beneficial (or less pathogenic). Second, drought increased the number of ears produced per plant, and more ears are produced with drought-conditioned microbes (Fig. 2H); however, multiple ear production is often a symptom of stress or poor pollination. Thus, plants producing more ears under drought when treated with drought-conditioned microbes is also likely consistent with microbe-mediated mal-acclimation rather than acclimation, particularly as we observed patterns of microbe-mediated mal-acclimation in kernel production (Fig. 2FG).

**Fig. 2.**
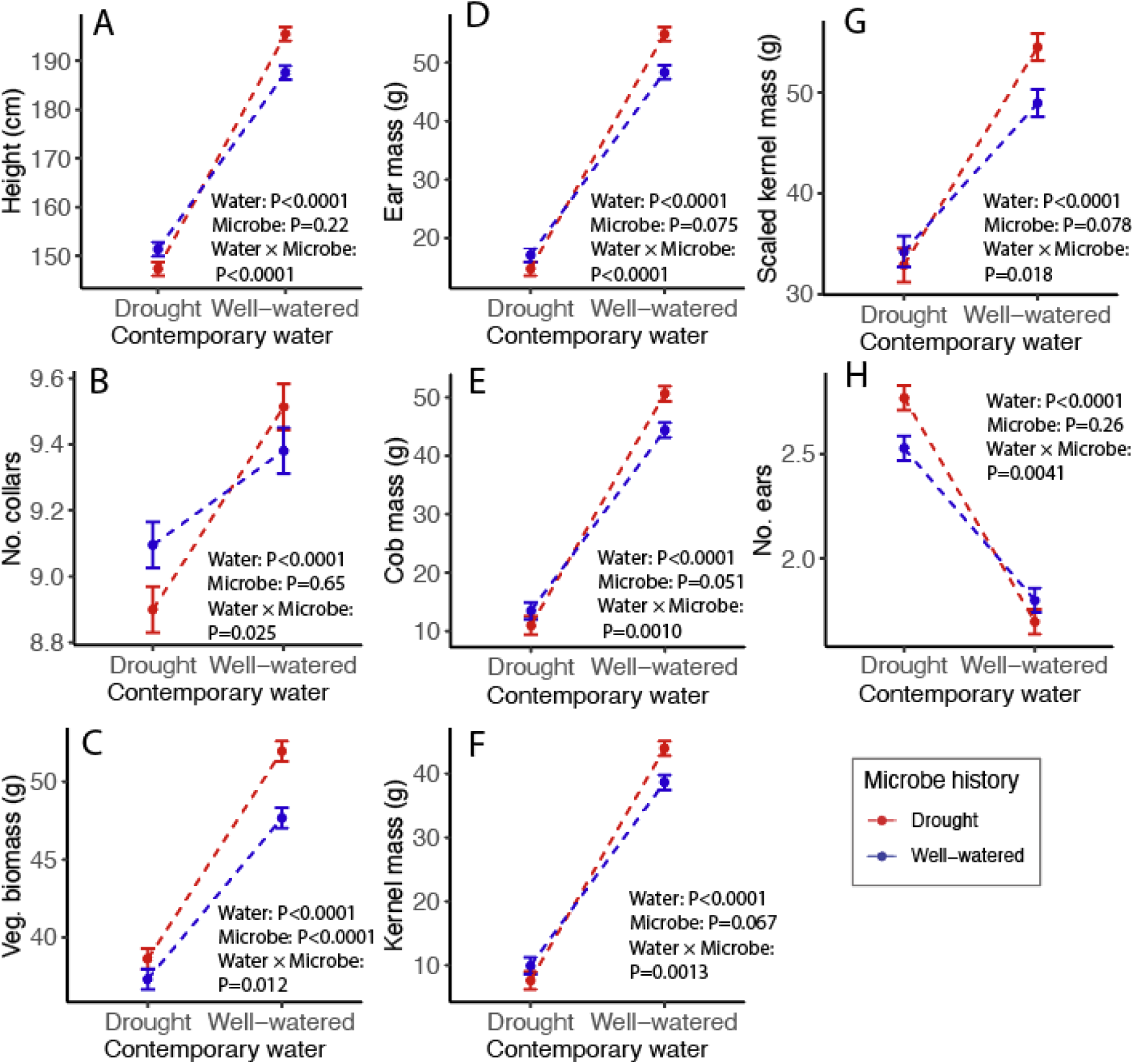
In general, we observed patterns consistent with microbe-mediated mal-acclimation, where performance of plants inoculated with microbes from drought conditions exacerbated the effects of drought stress on plant growth and reproduction. Soil microbial communities originated from 21 maize farms treated with drought or well-watered conditions in a greenhouse with a maize plant (“microbe history”) and then inoculated onto plants growing under different watering treatments (“contemporary water”). Values shown are estimated marginal mean vegetative (A-C) and reproductive (D-H) plant performance metrics (± SE) averaged over inoculant source farm and conditioning replicate. P-values were adjusted with an FDR correction.

### Do farms vary in their capacity for microbe-mediated acclimation?

Microbial effects on plant acclimation to drought varied by farm inoculant source, as indicated by a three-way interaction between contemporary water treatment, microbe history, and farm source for plant height (^2^(1)=34.47, FDR-corrected P = 0.069; raw P = 0.023), but not for other plant performance metrics (all P >> 0.05) (Appendix S1: Table S2). Plant height is likely the best metric of plant performance as it was measured during the middle of the growth period and was thus not limited by pot size, a spider mite outbreak that may have limited later season growth, or pollination efficiency, which could have limited response variables measured at harvest. Based on plant height, the soils from one third of farms showed patterns consistent with microbe-mediated mal-acclimation (farms 5, 16, 28, 47 {marginally significant, P = 0.053}, 51, 62, 63), whereas the inocula from the remaining farms did not respond to drought in ways that significantly affected plant responses to subsequent soil moisture conditions (Fig 3, Appendix S1: Table S3). While soils from some of the farms exhibited microbe-mediated mal-acclimation under both drought and well-watered conditions (farms 16, 47, and 51), microbe-mediated mal-acclimation was often only observed in one soil moisture environment. The soils from four farms showed mal-acclimation under well-watered conditions (farms 5, 28, 62, and 63) but not drought. In other words, microbes from wet environments inhibited plant growth in wet environments. While one farm (farm 49) did not exhibit significant microbe-mediated responses to soil moisture (nonsignificant contemporary water × microbe history interaction), it showed a trend of mal-acclimation under drought conditions. Thus, while we did not observe microbe-mediated acclimation or mal-acclimation in most (67 %) of the soil communities tested, microbe-mediated responses varied and microbe-mediated mal-acclimation was more common than acclimation (33% vs. 0%, respectively).

**Fig. 3.**
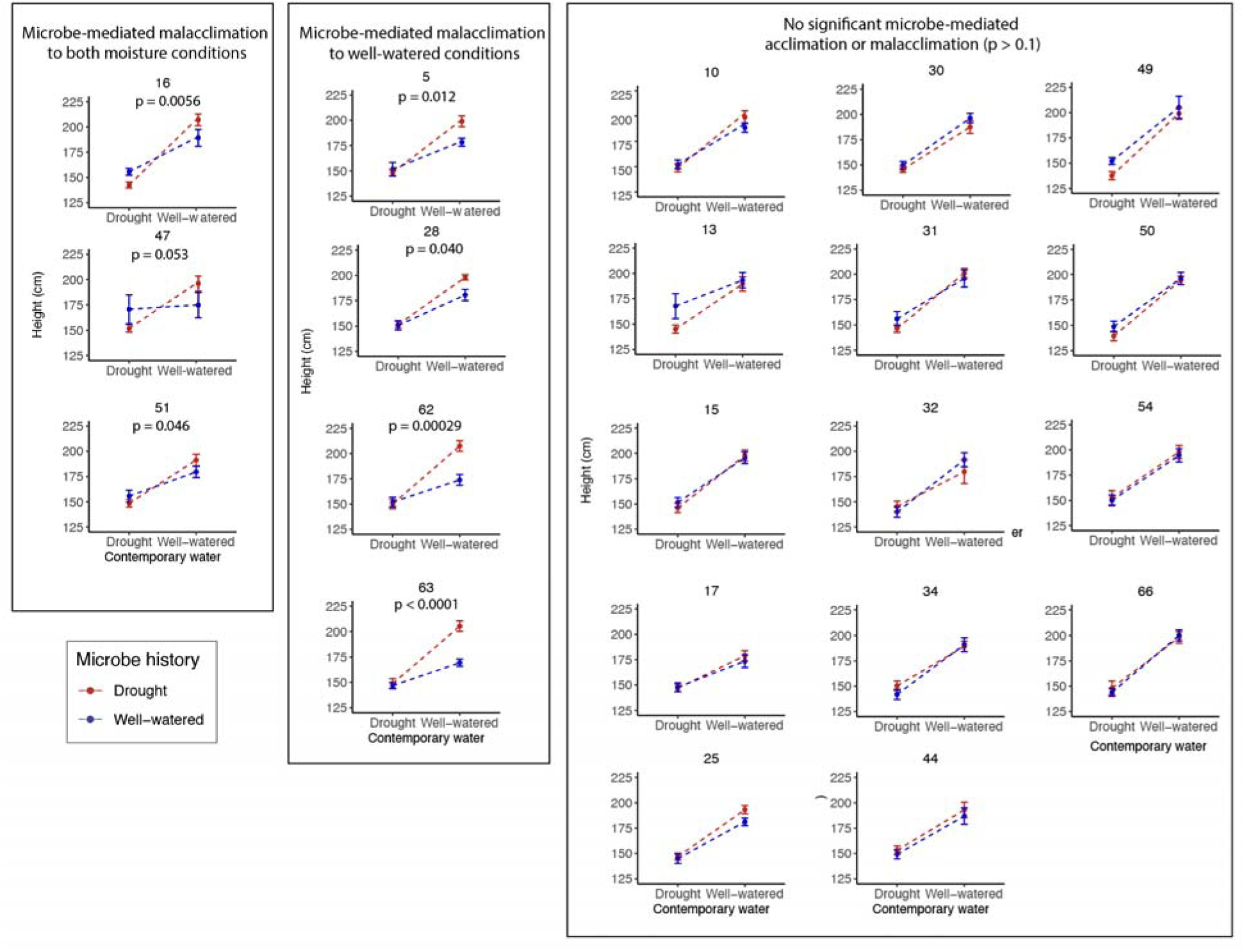
Variation in microbe-mediated plant acclimation to soil moisture across midwestern maize farms. Soil microbial communities from 21 maize farms were conditioned under drought or well-watered environments in a greenhouse with a maize plant (“microbe history”) and then inoculated onto plants growing under drought and well-watered contemporary environments (“contemporary water”). Soils from some farms caused plant mal-acclimation to both soil moisture conditions (left panel), while some only caused mal-acclimation in well-watered conditions (middle panel). Soils from most farms did not significantly affect plant acclimation to either soil moisture conditions (right panel). Values are estimated marginal mean plant heights (± SE) averaged over inoculant conditioning replicates. Numbers above each plot represent farm ID number. P-values indicate significant interactive effects between microbe history and contemporary water on plant height (see complete statistical results in Table S3).

We attempted to identify drivers of microbe-mediated acclimation/mal-acclimation through associations with soil and microbial properties. One of the PCs of the microbial properties marginally predicted microbe-mediated acclimation (FDR-corrected P=0.051, raw P=0.013), but the soil property PCs did not. The microbial community properties associated with the most predictive PC axis (PC2) included bacterial diversity, and the relative abundance of fungal root pathogens, dung saprotrophs, algal parasites, and unspecified saprotrophs. We also individually tested soil and microbial properties (microbial biomass, bacterial and fungal diversity, fungal pathogen relative abundance, mycorrhizal relative abundance, soil pH, soil texture), management practices (irrigation, tillage), and climate variables (mean annual precipitation and temperature) but none were significant predictors of microbe-mediated acclimation (Appendix S1: Table S7). However, the microbial communities that generated microbe-mediated mal-acclimation were all from soils that had low wet aggregate stability (<21%)(Appendix S1: Fig. S1), i.e., soils with structure that readily erodes upon wetting. However, wet aggregate stability did not significantly predict microbe-mediated (mal)acclimation (β_microbe history× contemporary water_) after correcting for testing multiple farm characteristics (FDR corrected P=0.593, raw P=0.0456).

## DISCUSSION

Understanding how and when microbes help or harm their hosts is an important area of both basic and applied research. While there are many examples of soil microbes helping plants acclimate to stressful conditions (Liu et al. 2020) and previous studies repeatedly find microbe-mediated acclimation to drought (Table 1), none of the soil microbial communities we tested helped maize plants acclimate to low soil moisture conditions. In fact, we sometimes observed the opposite phenomenon: microbe-mediated mal-acclimation. That is, microbes reduced the performance of plants under the soil moisture levels they had previously experienced. These microbe-mediated acclimative effects also varied widely between the microbial communities tested, with inocula from one third of farms showing mal-acclimative effects, while the remaining soil microbial communities did not significantly affect plant acclimation to soil moisture. These results demonstrate that the exposure of microbes to an environmental stress does not always improve–and may often impede–plants’ acclimation to stressors, raising questions about how and when microbe-mediated acclimation and mal-acclimation occur, and the consequences of mal-acclimative microbes.

### Mechanisms of microbe-mediated mal-acclimation

While microbe-mediated acclimation results from stressed-exposed microbial communities affecting plant physiology or modifying the abiotic environment, we believe microbe-mediated mal-acclimation likely stems from the deleterious effects of pathogens. Pathogens can rapidly adjust to changes in their environment such as water availability through shifts in abundance, community composition, and (likely to a lesser degree here) natural selection (Chakraborty 2013). We more commonly observed microbe-mediated mal-acclimation to wet soil rather than to drought. While wet soils may favor different pathogens than dry soils, pathogens generally accumulate in wetter soils (Cook and Papendick 1972). Thus, the accumulation of pathogens in soils conditioned under ample moisture, further amplified by high moisture in the response phase, likely reduced plant performance, resulting in microbe-mediated mal-acclimation. We also, albeit more rarely, observed microbe-mediated mal-acclimation to drought, which might result from selection for drought-tolerant pathogens that then imposed stronger negative effects on plant growth in the subsequent drought conditions. Interestingly, the initial relative abundance of putative pathogens in our farm soils was not significantly associated with microbe-mediated mal-acclimation in our study (p = 0.708), indicating that pathogen-laden soils are not necessarily prone to microbe-mediated mal-acclimation. However, it is possible that pathogen loads changed substantially in the greenhouse environment throughout the experiment (Hu et al. 2025).

Aside from affecting pathogens, drought may also cause declines in the abundance of beneficial microbes. Arbuscular mycorrhizal fungi, which can improve plant tolerance to drought through improving access to water and nutrients, often decline in abundance under drought (Gehring et al. 2017). This drought-induced reduction in beneficial fungi could explain why we sometimes observed plants growing more slowly under drought with drought-conditioned microbiomes. We did not find any significant associations between the initial relative abundance of mycorrhizal fungi in farm soils and microbe-mediated acclimation responses, however it is possible that there were low abundances of mycorrhizal fungi in our greenhouse experiment and the fungal amplicon sequencing approach that we used to characterize fungal communities in the field soils was not optimized for characterizing mycorrhizal fungi (Li et al. 2020).

### Is microbe-mediated acclimation less likely in agricultural systems?

While past studies documenting microbe-mediated acclimation to drought might lead one to conclude that it is common, none of the 21 soil communities we tested improved plant acclimation. There are several plausible explanations for this discrepancy. First, microbe-mediated acclimation may indeed be rare, and publication bias artificially inflates the perceived frequency of microbe-mediated acclimation (as opposed to mal-acclimation) in nature. Second, many studies documenting microbe-mediated acclimation compare soils across natural precipitation gradients or from multi-generation drought experiments. Perhaps soil microbial communities simply cannot respond strongly enough to the shorter (single growing season) drought imposed in our study, although this seems less likely given that we did detect small but statistically significant effects of microbe history (either main effects or interactive effects with contemporary moisture) in many of our soil communities. Third, microbe-mediated acclimation may be more common in natural ecosystems than agricultural ecosystems. Most studies reporting microbe-mediated acclimation examined microbial communities from natural grasslands or forest systems, while we focused on microbes from agricultural soils. Conventionally-managed agricultural soils are subject to many perturbations including fertilizers, pesticides, and tillage, and typically harbor a higher abundance of pathogens (Ohigashi et al. 2025), which may promote microbe-mediated mal-acclimation (see above), especially given that the homogeneity of industrial farms also favors the rapid evolution of virulence and host specialization in pathogens (Holguin and Bashan 1992). Additionally, soil microbial communities from grasslands are functionally more resilient to climate change (e.g., extreme hot/dry summers) than those from croplands (Bei et al. 2023), which suggests that natural landscapes may harbor communities with greater potential to adapt to stressful environmental conditions, which could translate into microbe-mediated acclimation.

In addition to differences in soil microbial communities, agricultural systems also contain very different plant hosts. Different plant species and genotypes vary in their drought tolerance and the degree to which they interact with microbes (dependence on microbes for stress tolerance, and susceptibility to pathogens), including under drought stress (Fitzpatrick et al. 2018). Domestication and breeding in resource-rich soils may have inadvertently reduced selection for traits that promote beneficial plant-microbe interactions (Pérez-Jaramillo et al. 2018; Porter and Sachs 2020), potentially making microbe-mediated acclimation more likely in wild species than modern crops. Furthermore, planting new seeds each year prevents the local adaptation of host plants to the environment. Additionally, several of the documented examples of microbe-mediated acclimation to drought were found in cruciferous species (Table 1), which may be less susceptible to pathogens due to their production of antimicrobial glucosinolates (Poveda et al. 2020). Such pathogen resistance would reduce the deleterious effects of acclimated pathogens that likely underlie maladaptive effects. Likewise, the unique chemistry of maize, particularly under drought stress, may affect its susceptibility to pathogens and its interactions with soil microbes. During drought, maize plants produce antimicrobial benzoxazinoids (Richardson and Bacon 1993), which are often released into the soil via root exudates, altering the plants’ rhizosphere communities (Hu et al. 2018). Benzoxazinoids can so strongly alter microbial interactions in the rhizosphere that they effectively mitigate negative plant-soil feedbacks (Bass 2024; Gfeller et al. 2024). The increased production of benzoxazinoids under drought may explain why we rarely observed microbe-mediated mal-acclimative responses in dry conditions (high benzoxazinoid levels kill rhizosphere pathogens), but observed them frequently in well-watered conditions, where low titers of benzoxazinoids likely are not able to combat the high pathogen loads of moist soils. In sum, plant physiological traits likely influence the likelihood of microbes substantially affecting plant acclimation to stressful conditions, and our focus on a modern maize variety may have made us less likely to observe microbe-mediated acclimation and more likely to observe mal-acclimation to drought.

### Conclusions and implications of microbe-mediated mal-acclimation

In sum, while there are many examples of microbe-mediated acclimation to drought in natural systems, to the best of our knowledge, microbe-mediated stress acclimation has not been studied in agricultural ecosystems. We did not find any evidence for this phenomenon in soil communities from large-scale maize farms from three midwestern states. Instead, we observed a general effect of microbe-mediated *mal-acclimation* to soil moisture conditions, with the effects varying widely between different source microbial communities. The soil communities from six of the farms exhibited statistically significant microbe-mediated mal-acclimation under at least one soil moisture environment (plus one marginally significant farm). While few individual soil, management, or microbial properties were predictive of microbe-mediated (mal)acclimation, our data suggest soil microbial communities from soils with low wet aggregate stability and microbial communities characterized by high bacterial diversity and relative abundances of fungal root pathogens, dung saprotrophs, algal parasites, and unspecified saprotrophs may be more likely to harbor microbial communities that impair acclimation to soil moisture conditions. Though neither tillage or irrigation influenced microbe-mediated acclimation in our study, identifying management practices that cultivate microbial communities that improve the capacity for microbe-mediated acclimation, or at least minimize microbe-mediated mal-acclimation could enhance the resilience of agricultural landscapes. Such management may become increasingly important because if microbe-mediated mal-acclimation is common, as our results suggest, the effects of extreme droughts (or deluges) on plants will worsen if these events become more frequent and likely to occur in consecutive growing seasons. On the other hand, many climate models predict increasing variability in precipitation (Pendergrass et al. 2017), which could benefit plant performance in cases where microbes cause mal-acclimation, as wet conditions would follow dry conditions and vice versa. Thus, understanding how and when microbes influence plant acclimation is important for predicting plant productivity and the stability of crop yields. As soil health increasingly becomes a priority for farmers (Irvine et al. 2023), our findings raise questions about the properties of soil communities that support plant resilience and how we could promote these beneficial soils in agriculture.

## ACKNOWLEDGMENTS

This work was supported by National Science Foundation Award BCS-2009125 to JAL, JTL, SE, SMP, by the NSF Long-term Ecological Research Program (DEB 2224712) at the Kellogg Biological Station, and by Michigan State University AgBioResearch. We also thank Maddie Gellinger, Jordan Ziss, Caleb Callahan, Kim Reish, and Jason Winters for assistance with the greenhouse experiment, Kristina Beethem for assistance with identifying management properties of the studied farms, Matt Houser and Rachel Irvine for establishing connections with the farms from which collected soil, Jipeng Luo for helpful discussions and feedback on this manuscript, and especially the farmers from Indiana, Michigan, and Illinois who allowed us to access their fields and sample their soils.

## AUTHOR CONTRIBUTIONS

**Mia Howard:** Writing – original draft, Formal analysis, Data curation, Investigation. **Lana Bolin:** Writing – review and editing, Formal analysis, Data curation, Conceptualization, Investigation. **Glade Bogar:** Writing – review and editing, Data curation, Investigation. **Sarah Evans:** Writing – review and editing, Conceptualization, Funding acquisition**. Jay Lennon:** Writing – review and editing, Conceptualization, Funding acquisition. **Sandy Marquart-Pyatt:** Writing – review and editing, Conceptualization, Funding acquisition, Data curation, Investigation. **Jennifer Lau:** Writing – review and editing, Conceptualization, Funding acquisition, Data curation, Investigation.

