## Supplemental Materials for "Microbe-mediated plant acclimation to drought may be rare in agriculture"

APPENDIX S1.

Title: Microbe-mediated plant acclimation to drought may be rare in agriculture

Authors: Mia M. Howard, Lana G. Bolin, Glade D. Bogar, Sarah E. Evans, Jay T. Lennon, Sandra T. Marquart-Pyatt, Jennifer A. Lau

Section S1: Methods

*Soil physical and chemical analyses*

We calculated soil moisture, field-moist, air-dry, and oven-dry conversion factors by weighing a subsample of field-moist soil onto aluminum dishes, weighing it after air drying it for 72 h, and finally weighing it after drying it in an oven at 105°C overnight. We calculated average bulk density by multiplying total sampled volume by total moisture-corrected soil mass. We measured soil pH by suspending 5.0 g field-moist soil in 10.0 mL 0.01 M CaCl2, incubating 30 min at room temperature, briefly centrifuging, and measuring with a FiveEasy Plus pH meter (Mettler Toledo). Other soil chemical and physical analyses were performed on air-dry soils at Brookside Laboratories, Inc. (New Bremen, OH) including soil texture (Bouyoucos, 1962), organic matter (Schulte and Hopkins, 1996), Total %C, %N, and C/N ratio (Nelson and Sommers, 1996), Bray 1 Phosphorus (Bray and Kurtz, 1945), cation exchange capacity (Ross and Ketterings, 1995), and Mehlich III Extractable P, Mn, Zn, B, Cu, Fe, Al, S, Ca Mg, K, and Na (Mehlich, 1984). Soil texture varied, with soils ranging from 4 to 78% sand and most soils classified as sandy loam or silt loam.

*Soil biochemical analyses*

Additional biochemical analyses were conducted to suggest potential mechanisms for microbe-mediated acclimation to drought: biomass, extracellular polymeric substance (EPS) content, and soil wet aggregate stability. Microbial biomass was measured using a chloroform fumigation method in which 2.0 mL of chloroform was added to 6.0 g of triplicate field-moist soil in 50 mL falcon tubes, capped for 24 hours then vented for 24 hours prior to extraction with 0.5M K_2_SO_4_ (Vance et al. 1987). EPS was extracted and quantified as described in (Bogar et al. 2025). Wet Aggregate Stability was quantified at the Cornell Soil Health Laboratory (Ithaca, NY).

*Soil microbial communities*

DNA was extracted from frozen subsamples and negative (no sample) controls using Qiagen DNeasy PowerSoil Pro kits (Qiagen, Carlsbad, CA). Extraction yield was quantified using a Qubit Fluorometer (Thermo Fisher, Waltham, MA). Bacterial amplicons were generated at the Michigan State University Research Technology Support Facility (RTSF) Genome Core. Bacterial library preparation was conducted by amplifying the V4 region of 16S genes (515f/806r, (Kozich et al. 2013)), using PCR with Dream Taq Master Mix (K9012) and the following cycling conditions: 95°C for 3 min, 30 cycles of 95°C for 45 s/50°C for 60 s/72°C for 90 s , and a final extension step at 72°C for 10 min. Fungal amplicons were generated using a modified version of the Earth Microbiome Project ITS amplification protocol (Bokulich and Mills 2013). Briefly, ITS genes (ITS1f/ITS2, (Smith and Peay 2014)) were amplified using PCR with Platinum II Hot-Start PCR Master Mix and the following cycling conditions: 94°C for 1 min, 25 cycles of 94°C for 30 s/52°C for 30 s/68°C for 30 s, and a final extension step at 68°C for 10 min. The cycle number was chosen to be as low as possible to minimize PCR bias (Chen et al. 2021) while allowing products to be visualized on a gel. ITS amplicons were cleaned using Beckman AMPure beads (1:1 ratio). Barcoding, quality control, normalization, pooling, and sequencing was performed by the MSU RTSF Genome Core, according to their protocols. Sequencing was performed on an Illumina MiSeq.

Amplicon reads were quality-screened and trimmed of primers using cutadapt (version 4.2 using Python v3.10.8; Martin, 2011), and bioinformatic analyses were performed using DADA2 (Callahan et al. 2016) using default settings, except as follows. For ITS, reads were quality-filtered using filterandTrim() with minLen = 50. For 16S, reads were quality-filtered using filterandTrim() with maxEE = c(5, 5), truncLen=c(220,180), and minLen = 100.Taxonomy was assigned to quality-filtered, denoised, merged, and chimera-filtered amplicon sequence variants (ASVs) using assignTaxonomy(). For ITS, we used UNITE reference “sh_general_release_dynamic_25.07.2023.fasta” (Nilsson et al. 2019); for 16S, SILVA reference “silva_nr99_v138.1_wSpecies_train_set.fa.gz” (Quast et al. 2013). Final read depth for 16S communities ranged from 6,099 - 24,934 (mean: 14,408); ITS communities ranged from 2,823 - 171,709 (mean 47,243). Rarefaction curves plateaued for all samples, suggesting adequate sequencing depth.

Communities along with their environmental metadata were compiled into phyloseq objects (McMurdie and Holmes 2013) for further analysis. Fungal traits and niches were assigned to ITS ASVs by referencing the FungalTraits database (Põlme et al. 2020). Shannon diversity was calculated using vegan (Oksanen et al., 2022).

Põlme, S., Abarenkov, K., Henrik Nilsson, R., Lindahl, B.D., Clemmensen, K.E., Kauserud, H., Nguyen, N., Kjøller, R., Bates, S.T., Baldrian, P., Frøslev, T.G., Adojaan, K., Vizzini, A., Suija, A., Pfister, D., Baral, H.-O., Järv, H., Madrid, H., Nordén, J., Liu, J.-K., Pawlowska, J., Põldmaa, K., Pärtel, K., Runnel, K., Hansen, K., Larsson, K.-H., Hyde, K.D., Sandoval-Denis, M., Smith, M.E., Toome-Heller, M., Wijayawardene, N.N., Menolli, N., Reynolds, N.K., Drenkhan, R., Maharachchikumbura, S.S.N., Gibertoni, T.B., Læssøe, T., Davis, W., Tokarev, Y., Corrales, A., Soares, A.M., Agan, A., Machado, A.R., Argüelles-Moyao, A., Detheridge, A., de Meiras-Ottoni, A., Verbeken, A., Dutta, A.K., Cui, B.-K., Pradeep, C.K., Marín, C., Stanton, D., Gohar, D., Wanasinghe, D.N., Otsing, E., Aslani, F., Griffith, G.W., Lumbsch, T.H., Grossart, H.-P., Masigol, H., Timling, I., Hiiesalu, I., Oja, J., Kupagme, J.Y., Geml, J., Alvarez-Manjarrez, J., Ilves, K., Loit, K., Adamson, K., Nara, K., Küngas, K., Rojas-Jimenez, K., Bitenieks, K., Irinyi, L., Nagy, L.G., Soonvald, L., Zhou, L.-W., Wagner, L., Aime, M.C., Öpik, M., Mujica, M.I., Metsoja, M., Ryberg, M., Vasar, M., Murata, M., Nelsen, M.P., Cleary, M., Samarakoon, M.C., Doilom, M., Bahram, M., Hagh-Doust, N., Dulya, O., Johnston, P., Kohout, P., Chen, Q., Tian, Q., Nandi, R., Amiri, R., Perera, R.H., dos Santos Chikowski, R., Mendes-Alvarenga, R.L., Garibay-Orijel, R., Gielen, R., Phookamsak, R., Jayawardena, R.S., Rahimlou, S., Karunarathna, S.C., Tibpromma, S., Brown, S.P., Sepp, S.-K., Mundra, S., Luo, Z.-H., Bose, T., Vahter, T., Netherway, T., Yang, T., May, T., Varga, T., Li, W., Coimbra, V.R.M., de Oliveira, V.R.T., de Lima, V.X., Mikryukov, V.S., Lu, Y., Matsuda, Y., Miyamoto, Y., Kõljalg, U., Tedersoo, L., 2020. FungalTraits: a user-friendly traits database of fungi and fungus-like stramenopiles. Fungal Divers. 105, 1–16. https://doi.org/10.1007/s13225-020-00466-2

Quast, C., Pruesse, E., Yilmaz, P., Gerken, J., Schweer, T., Yarza, P., Peplies, J., Glöckner, F.O., 2013. The SILVA ribosomal RNA gene database project: improved data processing and web-based tools. Nucleic Acids Res. 41, D590–D596. https://doi.org/10.1093/nar/gks1219

Ross, D.S., Ketterings, Q. 2011 Chapter 9 Recommended Methods for Determining Soil Cation Exchange Capacity. (NECC-1812), T.N.C.C.f.S.T. (ed).

Smith, D.P., Peay, K.G., 2014. Sequence Depth, Not PCR Replication, Improves Ecological Inference from Next Generation DNA Sequencing. PLOS ONE 9, e90234. https://doi.org/10.1371/journal.pone.0090234

Vance, E.D., Brookes, P.C., Jenkinson, D.S., 1987. An extraction method for measuring soil microbial biomass C. Soil Biol. Biochem. 19, 703–707. https://doi.org/10.1016/0038-0717(87)90052-6

Section S2: Supporting tables and figures

Table S1. Results of ANOVAs comparing the effects of contemporary watering environment and microbe history on the drought responses of maize plants. Symbols after the 𝛘^2^ value indicated statistical significance after correction with a FDR correction: *** <0.001, ** <0.01, * <0.05, ・<0.10.

|  | **Height** | **Number of collared leaves** | **Stem diameter** | **Vegetative biomass** | **Ear mass** | **Number of ears** | **Cob mass** | **Kernel mass** | **Scaled kernel mass** |
| --- | --- | --- | --- | --- | --- | --- | --- | --- | --- |
| Contemporary water | 𝛘^2^ = 1043.9*** | 𝛘^2^ = 42.2*** | 𝛘^2^ = 63.8*** | 𝛘^2^ = 404.9*** | 𝛘^2^ = 1192.2*** | 𝛘^2^ = 65.9*** | 𝛘^2^ = 786.0*** | 𝛘^2^ = 812.3*** | 𝛘^2^ = 172.1*** |
| Microbe history | 𝛘^2^ = 1.7 | 𝛘^2^ = 0.21 | 𝛘^2^ = 29.4*** | 𝛘^2^ = 20.2*** | 𝛘^2^ = 3.5・ | 𝛘^2^ = 0.40 | 𝛘^2^ = 4.3* | 𝛘^2^ = 3.8・ | 𝛘^2^ = 3.4・ |
| Contemporary water ✕ microbe  history | 𝛘^2^ = 20.9*** | 𝛘^2^ = 5.6* | 𝛘^2^ = 0.9 | 𝛘^2^ = 6.4* | 𝛘^2^ = 18.5*** | 𝛘^2^ = 2.2* | 𝛘^2^ = 12.1** | 𝛘^2^ = 11.4** | 𝛘^2^ = 6.3* |

Table S2. Results of ANOVAs comparing the effects of contemporary watering environment, microbe history, and soil origin (farm) on the drought responses of maize plants. A significant contemporary water ✕ microbe history ✕ farm interaction indicates microbe-mediated plant responses to soil moisture vary significantly by farm (inoculant source). Symbols after the 𝛘^2^ value indicated statistical significance after adjustment with a FDR: *** <0.001, ** <0.01, * <0.05, ・<0.10.

|  | **Height** | **Number of collared leaves** | **Stem diameter** | **Vegetative biomass** | **Ear mass** | **Number of ears** | **Cob mass** | **Kernel mass** | **Scaled kernel mass** |
| --- | --- | --- | --- | --- | --- | --- | --- | --- | --- |
| Contemporary water | 𝛘^2^= 1095.77*** | 𝛘^2^= 43.3*** | 𝛘^2^= 64.7*** | 𝛘^2^= 408.6*** | 𝛘^2^= 1207.1*** | 𝛘^2^=64.4*** | 𝛘^2^= 777.4*** | 𝛘^2^= 806.1*** | 𝛘^2^= 176.3*** |
| Microbe history | 𝛘^2^= 1.8 | 𝛘^2^= 0.2 | 𝛘^2^=29.2*** | 𝛘^2^= 22.2*** | 𝛘^2^= 3.3 | 𝛘^2^= 0.4 | 𝛘^2^= 4.0 | 𝛘^2^= 3.5 | 𝛘^2^= 3.3 |
| Farm | 𝛘^2^= 21.9 | 𝛘^2^= 16.9 | 𝛘^2^= 25.4 | 𝛘^2^= 35.6* | 𝛘^2^= 31.7 | 𝛘^2^=2.6 | 𝛘^2^= 31.0 | 𝛘^2^= 31.0 | 𝛘^2^= 24.3 |
| Contemporary water ✕ microbe history | 𝛘^2^= 21.9*** | 𝛘^2^= 5.8* | 𝛘^2^= 0.9 | 𝛘^2^= 6.6* | 𝛘^2^= 18.8*** | 𝛘^2^= 2.0 | 𝛘^2^= 13.2*** | 𝛘^2^= 12.5** | 𝛘^2^= 8.8* |
| Contemporary water ✕ farm | 𝛘^2^= 36.7* | 𝛘^2^= 27.5 | 𝛘^2^= 31.2 | 𝛘^2^= 25.4 | 𝛘^2^= 33.1ç | 𝛘^2^= 2.7 | 𝛘^2^= 18.7 | 𝛘^2^= 19.8 | 𝛘^2^= 17.1 |
| Microbe history ✕ farm | 𝛘^2^= 26.9 | 𝛘^2^= 31.0 | 𝛘^2^= 19.1 | 𝛘^2^= 29.4 | 𝛘^2^= 12.1 | 𝛘^2^= 6.6 | 𝛘^2^= 10.8 | 𝛘^2^= 10.7 | 𝛘^2^= 28.3 |
| Contemporary water ✕ microbe history ✕ farm | 𝛘^2^=34.5・ | 𝛘^2^= 26.3 | 𝛘^2^= 17.0 | 𝛘^2^= 20.1 | 𝛘^2^= 15.2 | 𝛘^2^= 8.9 | 𝛘^2^= 21.2 | 𝛘^2^= 22.3 | 𝛘^2^= 21.0 |

Table S3. Results of ANOVAs comparing the effects of contemporary watering environment and microbe history on the drought responses of maize plants across farms. The response variable is plant stem height.

| **Farm ID** | **Contemporary water** | **Microbe history** | **Contemporary water ✕ microbe history** |
| --- | --- | --- | --- |
| 5 | 𝛘^2^=65.1  P<0.0001 | 𝛘^2^= 2.1  P=0.15 | 𝛘^2^= 6.3  P=0.012 |
| 10 | 𝛘^2^= 90.4  P<0.0001 | 𝛘^2^= 0.5  P=0.47 | 𝛘^2^= 1.9  P=0.16 |
| 13 | 𝛘^2^= 23.0  P<0.0001 | 𝛘^2^= 1.0  P=0.31 | 𝛘^2^= 1.7  P=0.20 |
| 15 | 𝛘^2^= 87.0  P<0.0001 | 𝛘^2^= 0.10  P=0.75 | 𝛘^2^= 0.6  P=0.45 |
| 16 | 𝛘^2^= 77.6  P<0.0001 | 𝛘^2^= 0.18  P=0.67 | 𝛘^2^= 7.7  P=0.0056 |
| 17 | 𝛘^2^= 36.0  P<0.0001 | 𝛘^2^= 0.2  P=0.68 | 𝛘^2^= 0.4  P=0.53 |
| 25 | 𝛘^2^= 106.9  P<0.0001 | 𝛘^2^= 3.0  P=0.084 | 𝛘^2^= 1.5  P=0.21 |
| 28 | 𝛘^2^= 87.4  P<0.0001 | 𝛘^2^= 3.2  P=0.073 | 𝛘^2^= 4.2  P=0.040 |
| 30 | 𝛘^2^= 95.2  P<0.0001 | 𝛘^2^= 1.5  P=0.21 | 𝛘^2^= 0.3  P=0.61 |
| 31 | 𝛘^2^= 70.9  P<0.0001 | 𝛘^2^= 0.05  P=0.83 | 𝛘^2^= 1.8  P=0.19 |
| 32 | 𝛘^2^= 33.0  P<0.0001 | 𝛘^2^= 0.1  P=0.73 | 𝛘^2^= 1.3  P=0.25 |
| 34 | 𝛘^2^= 64.5  P<0.0001 | 𝛘^2^= 0.4  P=0.54 | 𝛘^2^= 0.9  P=0.35 |
| 44 | 𝛘^2^= 37.3  P<0.0001 | 𝛘^2^= 0.62  P=0.43 | 𝛘^2^= 0.03  P=0.87 |
| 47 | 𝛘^2^= 5.5  P=0.019 | 𝛘^2^= 0.01  P=0.91 | 𝛘^2^= 3.8  P=0.053 |
| 49 | 𝛘^2^= 78.5  P<0.0001 | 𝛘^2^= 1.4  P=0.23 | 𝛘^2^= 0.4  P=0.52 |
| 50 | 𝛘^2^= 114.9  P<0.0001 | 𝛘^2^= 0.6  P=0.44 | 𝛘^2^= 0.6  P=0.43 |
| 51 | 𝛘^2^= 50.0  P<0.0001 | 𝛘^2^= 0.08  P=0.77 | 𝛘^2^= 4.0  P=0.046 |
| 54 | 𝛘^2^= 49.3  P<0.0001 | 𝛘^2^= 0.2  P=0.64 | 𝛘^2^= 0.004  P=0.95 |
| 62 | 𝛘^2^= 65.8  P<0.0001 | 𝛘^2^= 7.1  P=0.0077 | 𝛘^2^= 13.1  P=0.00029 |
| 63 | 𝛘^2^= 104.1  P<0.0001 | 𝛘^2^= 9.9  P=0.0017 | 𝛘^2^= 19.0  P=<0.0001 |
| 66 | 𝛘^2^= 93.6  P<0.0001 | 𝛘^2^= 0.04  P=0.84 | 𝛘^2^= 0.33  P=0.56 |

Table S4. Soil and microbial properties as predictors of microbe mediated acclimation to soil moisture. Soil and microbial properties were compressed using PCA. The first two principal components of the soil (Table S6) and microbial properties (Table S7). PCAs were assessed as predictors of microbe-mediated acclimation (represented as the model coefficient of the microbe history*contemporary water interaction term for each farm from the analysis in Fig. 3 and Table S3)

| **Predictor** | ***F*** | ***P (raw)*** | ***P (corrected)*** |
| --- | --- | --- | --- |
| Soil PC1 | 0.0238 | 0.882 | 0.954 |
| Soil PC2 | 1.741 | 0.214 | 0.428 |
| Microbial PC1 | 0.0035 | 0.954 | 0.954 |
| Microbial PC2 | 8.843 | **0.0127** | 0.051 |

Table S5. Variable loadings from PCA of microbial properties.

| **Property** | **PC1 (30% of variance)** | **PC2 (17% of variance)** |
| --- | --- | --- |
| Bacterial diversity | -0.1219752 | -0.3486475 |
| Fungal diversity | 0.32617549 | -0.3078817 |
| Arbuscular mycorrhizal fungi | -0.5444018 | 0.23216564 |
| Dung saprotroph | -0.2604805 | -0.7826796 |
| Foliar endophyte | 0.46597627 | -0.4586688 |
| Litter saprotroph | -0.4121446 | -0.0999398 |
| Mycoparasite | 0.45426539 | 0.34239364 |
| Nectar tap saprotroph | 0.33372556 | -0.0155296 |
| Plant pathogen | 0.78071089 | -0.2283639 |
| Root endophyte | -0.4323159 | 0.34318628 |
| Soil saprotroph | -0.829739 | 0.04386659 |
| Unspecified saprotroph | -0.1521361 | 0.60541021 |
| Wood saprotroph | 0.57206321 | 0.12750124 |
| Algal parasite | -0.3257818 | -0.5555125 |
| Root associated | -0.7513144 | 0.28042744 |
| Root pathogen | -0.3097165 | -0.6535844 |
| Leaf/fruit/seed pathogen | 0.81003282 | -0.1819964 |
| Microbial biomass | 0.51644578 | 0.42965115 |
| Exopolysaccharides | 0.50726412 | 0.2713257 |

Table S6. Variable loadings from PCA of soil properties.

| **Property** | **PC1 (38% of variance)** | **PC2 (19% of variance)** |
| --- | --- | --- |
| Sand | -0.6708942 | 0.34910916 |
| Silt | 0.54182931 | -0.3524991 |
| Clay | 0.80223649 | -0.2021159 |
| Cation exchange capacity | 0.72785366 | -0.2920947 |
| pH | -0.2200729 | 0.52660272 |
| C | 0.83344049 | -0.0877024 |
| N | 0.76162386 | -0.2401667 |
| Organic matter | 0.79863458 | -0.2488458 |
| P (Bray) | 0.43564741 | 0.61432647 |
| S | 0.05980893 | 0.7201853 |
| P | 0.49545625 | 0.61229211 |
| Ca | 0.65685098 | -0.132544 |
| Mg | 0.6680701 | -0.0314669 |
| K | 0.66948603 | 0.29040986 |
| Na | 0.01330228 | 0.37798211 |
| B | 0.29012322 | 0.61272342 |
| Fe | 0.27537846 | 0.22373678 |
| Mn | -0.289923 | 0.31317376 |
| Cu | 0.36756622 | 0.20632396 |
| Zn | 0.5316676 | 0.6000111 |
| Wet aggregate stability | -0.3337114 | -0.3647869 |

Table S7. Microbe, soil, climate, and management characteristics as predictors of microbe-mediated acclimation. We tested whether each inoculant characteristic predicted the coefficient of the microbe history*contemporary water interaction term from the models of plant height for each farm (models in Fig. 3/Table S3). A significant p-value would indicate that the inoculant characteristic predicts the microbe-mediated acclimative response.

| **Predictor** | ***F*** | ***P (raw)*** | ***P (corrected)*** |
| --- | --- | --- | --- |
| Soil bacterial diversity | 1.435 | 0.942 | 0.942 |
| Soil fungal diversity | 0.7154 | 0.412 | 0.783 |
| Soil fungal plant pathogen abundance | 2.4523 | 0.134 | 0.605 |
| Soil fungal mycorrhizal abundance | 0.1444 | 0.708 | 0.795 |
| Soil pH | 0.2331 | 0.635 | 0.795 |
| Soil texture (% sand) | 0.2297 | 0.637 | 0.795 |
| Soil exopolysaccharides | 2.1302 | 0.161 | 0.605 |
| Soil microbial biomass | 1.880 | 0.186 | 0.605 |
| Soil wet aggregate stability | 4.5756 | **0.0456** | 0.593 |
| Farm irrigation (irrigated or non-irrigated) | 0.1191 | 0.734 | 0.795 |
| Farm tillage (no-till, conservation, or conventional) | 1.2536 | 0.309 | 0.783 |
| Mean annual precipitation | 0.5026 | 0.487 | 0.791 |
| Mean annual temperature | 0.6742 | 0.422 | 0.783 |

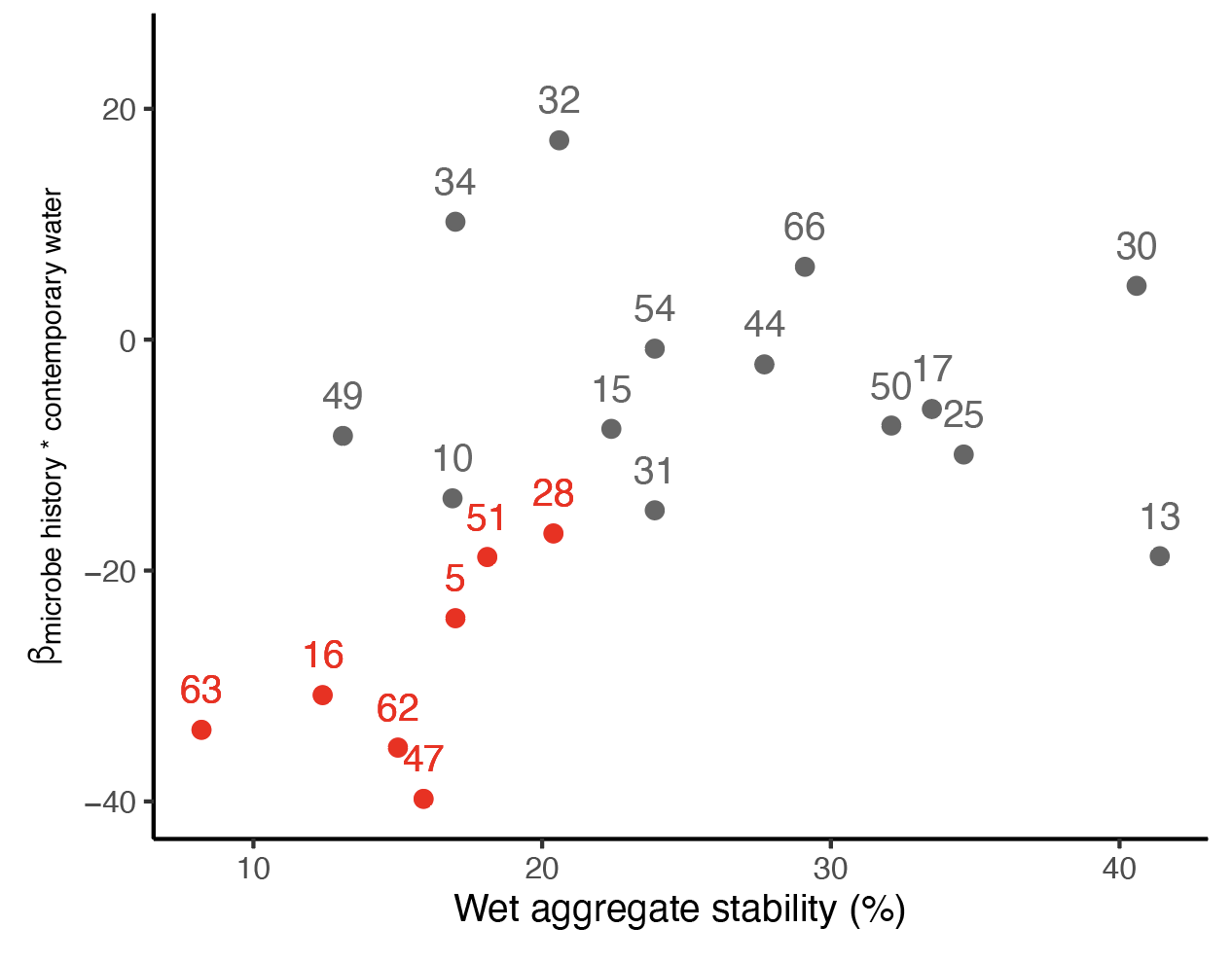

Fig. S1. Relationship between soil wet aggregate stability and microbe-mediated acclimation to soil moisture (represented as the model coefficient of the microbe history*contemporary water interaction term for each farm from the analysis in Fig. 3 and Table S3). Soils from farms that exhibited microbe-mediated mal-acclimation are colored in red.
